# Subjective rather than absolute reward value determines long-term memory formation in honey bees

**DOI:** 10.64898/2026.04.22.720144

**Authors:** A Charalambous, M Azcueta, R Barrozo, F Locatelli, M Klappenbach

## Abstract

How animals evaluate reward quality is a fundamental question in neuroscience and behavioral biology. Here we show that in honey bees (*Apis mellifera*), the value of a sucrose reward is not processed in absolute terms but relative to prior experience, and that this subjective evaluation strongly influences long-term memory formation. Using appetitive olfactory conditioning of the proboscis extension reflex (PER), we demonstrate that memory performance is determined by the contrast between a previously experienced reward and the reward used during training, rather than by the absolute concentration of sucrose received. This effect operates across multiple timescales, from contrasts between successive trials within a single session to differences between rewards experienced 24 hours apart. We further show that prior exposure to sucrose solutions of different concentrations modulates gustatory responsiveness and alters the sensitivity of antennal gustatory receptor neurons, suggesting that peripheral sensory plasticity contributes to experience-dependent changes in reward evaluation. Dissociating pre- and post-ingestive reward components revealed that the contrast between the sucrose concentration sensed by the antennae and the concentration ingested is sufficient to modulate memory formation. Together, our results indicate that bees form an internal expectation of reward quality based on experience, and that this expectation rescales the perceived value of subsequent rewards, thereby shaping associative memory strength. These findings provide a mechanistic framework for understanding how invertebrates perform relative reward comparisons across multiple temporal scales, with implications for flexible foraging strategies in dynamic environments.

## Introduction

How do animals evaluate reward quality, and how does experience bias their judgments of that reward?? The relative evaluation of reward quality is a common feature of human experience, whereby the perceived value of an outcome is determined not only by its absolute properties but also by expectations formed through prior experience. These questions were initially explored in mammals, but they can be addressed using a wide range of species. In a pioneering study, Tinklepaugh demonstrated how monkeys compare rewards through a simple experiment: he showed a banana to a monkey, but before the animal could reach it, he replaced it with lettuce. Under these circumstances, the monkey rejected the lettuce, despite normally accepting it as food (Tinklepaugh, 1928). This suggests that the animal was not evaluating the reward in absolute terms but rather comparing it with an expectation. Over the years, further evidence has been gathered using different behavioural paradigms and different vertebrate models. Research has shown that animals establish both positive and negative contrasts between past and present rewards, influencing their behaviour. The incentive contrast effect was first systematically studied in rats by L.P. Crespi and is therefore also known as the ‘Crespi effect’ (Crespi, 1942). However, less is known about how previous rewards influence the evaluation of a current reward in invertebrates (Couvillon and Bitterman, 1984; Hemingway and Muth, 2022; Wendt et al., 2019; Wiegmann et al., 2003).

Experiments based on foraging behaviour showed that fluctuations in food source profitability affect honeybees’ foraging preferences. Bees remember how reward quality changes over time: if they encounter a food source that provides increasing rewards, they visit it more persistently than another that offers decreasing rewards over time (Gil et al., 2007; Gil and De Marco, 2009). It was demonstrated that this schedule of rewards modifies the unconditioned response to a reward (Gil et al., 2008). Upon returning to the hive, forager bees recruit hive mates through various mechanisms, such as trophallaxis and the waggle dance, with the probability and intensity of recruitment being linked to the profitability of the food source (De Marco et al., 2005; Richter and Waddington, 1993). Using a sucrose responsiveness assessment, it has been demonstrated that experience alters how they perceive a sucrose reward solution. Bees that have previously encountered a more concentrated sugar solution respond less strongly in a subsequent test than those that experienced a less concentrated solution (Pankiw et al., 2001; Ramírez et al., 2010).

Although incentive contrast effects in honeybees have been characterized behaviourally, it remains unclear where along the reward-processing hierarchy these effects arise. Reward evaluation may be shaped by plasticity at multiple levels, ranging from peripheral sensory neurons to central circuits involved in learning and memory. Here, we combine both behavioural and physiological assays to investigate how experience modifies reward perception and how it impacts learning and memory in honeybees. Our findings show that prior experience modifies the response of gustatory receptor neurons, thereby contributing to changes in reward perception that strongly influence learning performance.

## Materials & Methods

### Animals

Pollen-foraging honeybees (*Apis mellifera*) were collected in the morning at the entrance of two regular hives located on the campus of the University of Buenos Aires (34°32′ S; 58°06′W). Bees were briefly cooled on ice and restrained in individual metal holders that allowed free movement of the antennae and proboscis. After recovery from cooling, animals were fed 1 µl of a sucrose solution and then remained undisturbed until being fed *ad libitum* in the evening. All sucrose solutions were prepared in distilled water.

In the laboratory, bees were kept in a humid chamber at room temperature (20–24 °C) under a 12:12 h light:dark cycle. All training and testing sessions were conducted between 10:00 and 14:00 during the austral spring–summer period.

### Olfactory conditioning

Honeybees were subjected to appetitive olfactory conditioning of the proboscis extension reflex (PER) following standard procedures (Bitterman et al., 1983). The odours used for conditioning were acetophenone and 1-hexanol, each diluted 1:10 in mineral oil (Sigma– Aldrich). For each experiment, 100 µl of the odour solution were freshly loaded into sealed 5 ml glass vials. Odour delivery was achieved via a constant airflow of charcoal-filtered air (500 ml/min) directed towards the bee’s head. During odour stimulation, a solenoid valve diverted a fraction of the airflow (50 ml/min) through the odour vial, carrying a portion of the headspace into the main airstream via a mixing chamber before reaching the animal. Thus, the final odour concentration reaching the animal was approximately one-tenth of the odour concentration in the vial headspace. The system was designed to maintain a constant final airflow, thereby preventing mechanical stimulation during odour presentation. A gentle air exhaust positioned 10 cm behind the bee continuously removed odours from the training arena.

During each training trial, an animal was placed in the training arena facing the odour delivery device. After 20 s, the odour was presented for 4 s. Three seconds after odour onset, the antennae were stimulated with a sucrose solution, the concentration of which varied depending on the experiment, eliciting the proboscis extension reflex. Upon proboscis extension, sucrose was delivered using a Gilmont micrometer syringe, allowing the bee to ingest 0.4 µl. Twenty seconds after reward consumption, the animal was returned to the resting position until the next trial. The intertrial interval was 10 min.

During training, responses were scored as positive when the proboscis was extended during odour presentation but prior to sucrose stimulation. The 3 s delay between odour onset and reward delivery allowed learning performance to be assessed during the training session. The same procedure was used during testing sessions, except that no reward was delivered.

### Sucrose responsiveness

Sucrose responsiveness was assessed following (R. Scheiner et al., 2001), with slight modifications. Bees were captured, immobilised, and fed as described previously. On the day of the experiment, bees were sequentially stimulated by touching the antennae with water followed by six sucrose solutions (0.1%, 0.3%, 1%, 3%, 10%, and 30% w/v). A sucrose responsiveness score was calculated as the total number of stimulations that elicited the PER, yielding scores ranging from 0 to 6. The inter-stimulus interval was 2 min.

### Recordings of antennal sensilla

Extracellular single-sensillum electrophysiology (Hodgson et al., 1955) was performed on chaetic sensilla located at the distal end of the antenna. The procedure was based on Haupt et al (Haupt, 2004), with slight modifications. Bees were captured and fed as described previously. To reduce electrical noise, bees were immobilised in plastic tips. On the following day, animals were chilled for 30 s to restrict antennal movement and fixed onto a microscope slide using adhesive tape. Animals were grounded via a silver (Ag/AgCl) wire inserted into the left optic lobe, which served as the reference electrode. A single sensillum was stimulated for 2 s using a glass capillary (20–30 µm tip diameter) containing a second silver (Ag/AgCl) wire acting as the recording electrode. The capillary was filled with the gustatory stimulus (sucrose at 0.1%, 1%, or 10% w/v) prepared in a 10 mM KCl electrolyte solution.

The recording electrode was connected to a preamplifier (Taste PROBE DTP-02, Syntech; gain ×10). Signals were subsequently amplified, filtered, and digitised using an amplifier (Dagan Ex1; gain ×100; band-pass filter: 1–3000 Hz; sampling rate: 10 kHz; 16-bit resolution) and an analogue-to-digital converter. Action potentials were identified and quantified using dbWave software (Marion-Poll, 1996).

### Statistical analysis

All statistical analyses are summarised in Table 1. Statistical analyses were conducted in R using generalized linear mixed models (GLMMs). The choice of model structure, error distribution, and link function was guided by the distributional properties of each response variable. Model diagnostics were assessed through inspection of residuals and fitted values (DHARMa). Statistical significance was evaluated using an alpha level of 0.05.

The proportion of Proboscis Extension Response (PER) was modeled using a binomial distribution with a logit link function. Experimental group and trial were included as fixed effects, and individual identity was included as a random intercept to account for repeated measurements. Training and test phases were modelled separately to distinguish two different processes (learning versus memory expression). For the training model, only trials 2 to 4 were analysed because animals that responded positively in the first trial were excluded; consequently, the response variable was equal to zero for all groups in trial 1. The models were fitted using the glmmTMB package in R. Post hoc pairwise comparisons between groups were performed using estimated marginal means with Tukey correction (Fig 2,3,5,6). For analyses focused on specific, predefined group comparisons, *a priori* orthogonal contrasts were used in figure 1,7,8.

**Figure 1.**
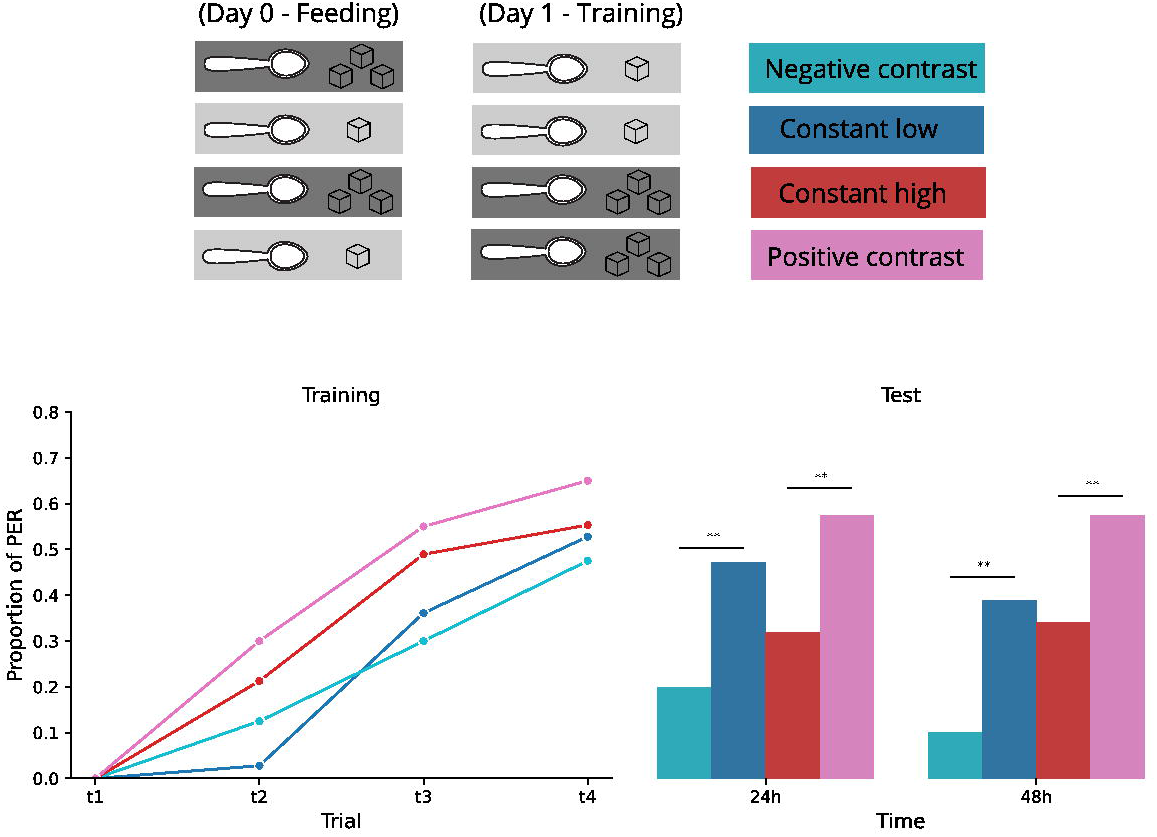
Feeding experience modulates memory strength. (a) Experimental protocol. On day 0, animals were fed with a sucrose solution of either 0.5 M (light grey) or 1.5 M (dark grey). On day 1, the same sucrose concentrations were used during a four trials training. The combination of feeding experience and reward during training defined four experimental groups: negative contrast (cyan, n = 40), constant low (blue, n =36), constant high (red, n = 47), and positive contrast (pink, n =40). (b) Left panel: line plots showing the proportion of positive responses for each group across the training trials. Right panel: responses during the test sessions 24 and 48 hours after training. **p < 0.01

**Figure 2.**
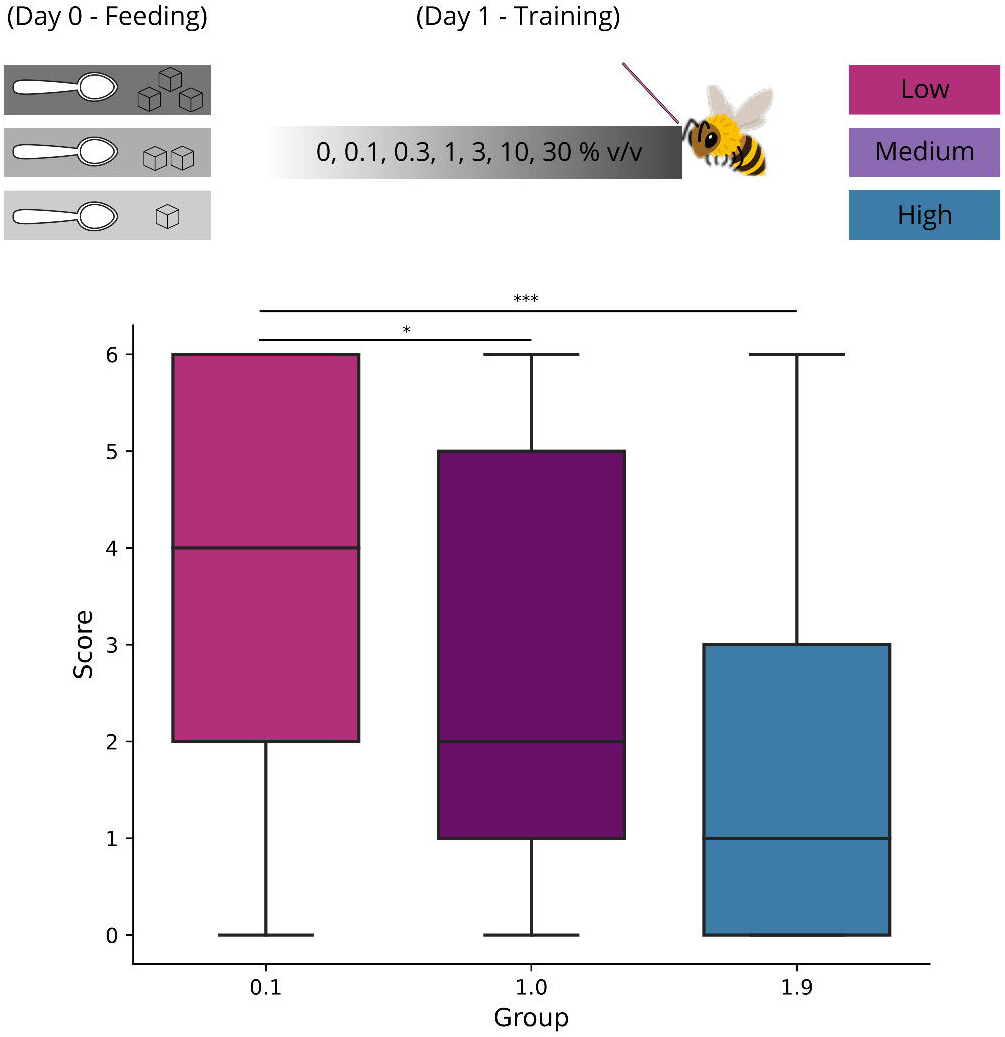
Feeding experience modulates sucrose responsiveness. (a) Experimental protocol. On day 0 animals were fed with a solution of sucrose 0.1M (light grey), 1M (medium grey) or 1.9M (dark grey) defining three experimental groups: low (magenta, n = 23), medium (violet, n = 39), and high (blue, n = 39). On day 1, all animals were stimulated with water and five solutions of increasing sucrose concentration. (b) Sucrose responsiveness score. The horizontal black line represents the median, boxes indicate the interquartile range, and whiskers denote the range. * p < 0.05, *** p < 0.001

**Figure 3:**
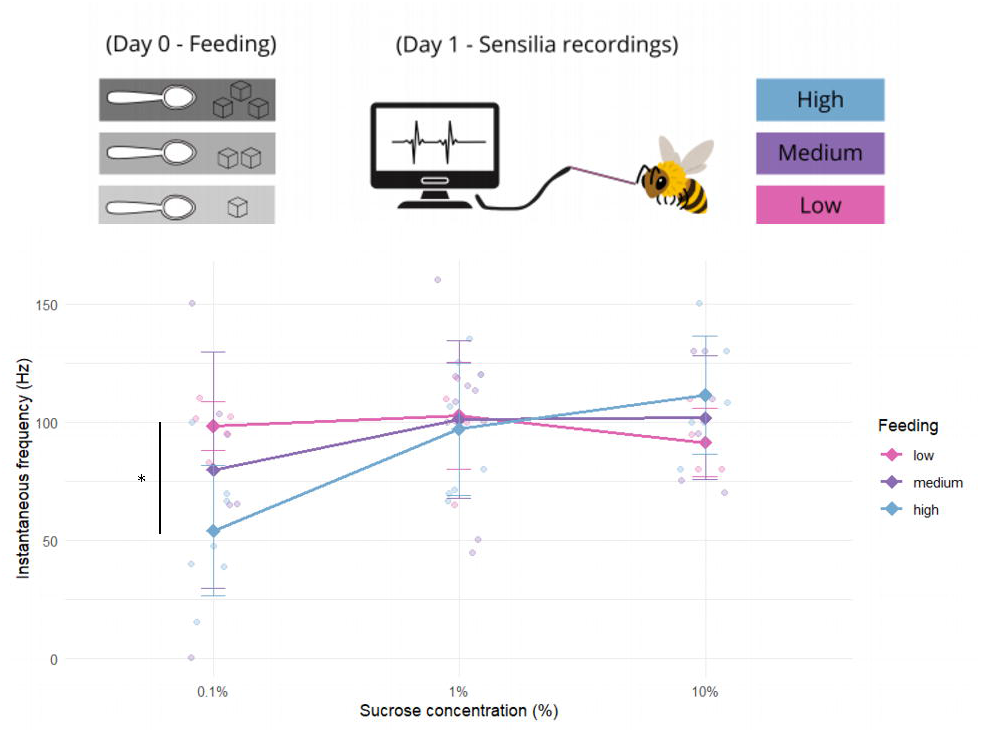
Antennal gustatory chemoreceptor response based on prior taste experience. Instantaneous frequency at 100 ms after stimulus onset for three experimental groups at three sucrose concentrations. Smaller points represent individual bees. Larger points represent the group means with their corresponding standard errors. Sample sizes: low - 0.1% (5bees, 9sensilia), medium - 0.1% (6bees, 12sensilia), high - 0.1% (7bees, 19 sensilia), low - 1% (5bees, 11sensilia), medium - 1% (10bees, 21sensilia), high - 1% (8bees, 13 sensilia), low - 10% (4bees, 5sensilia), medium - 10% (6bees, 7sensilia), high - 10% (6bees, 10 sensilia)

**Figure 4.**
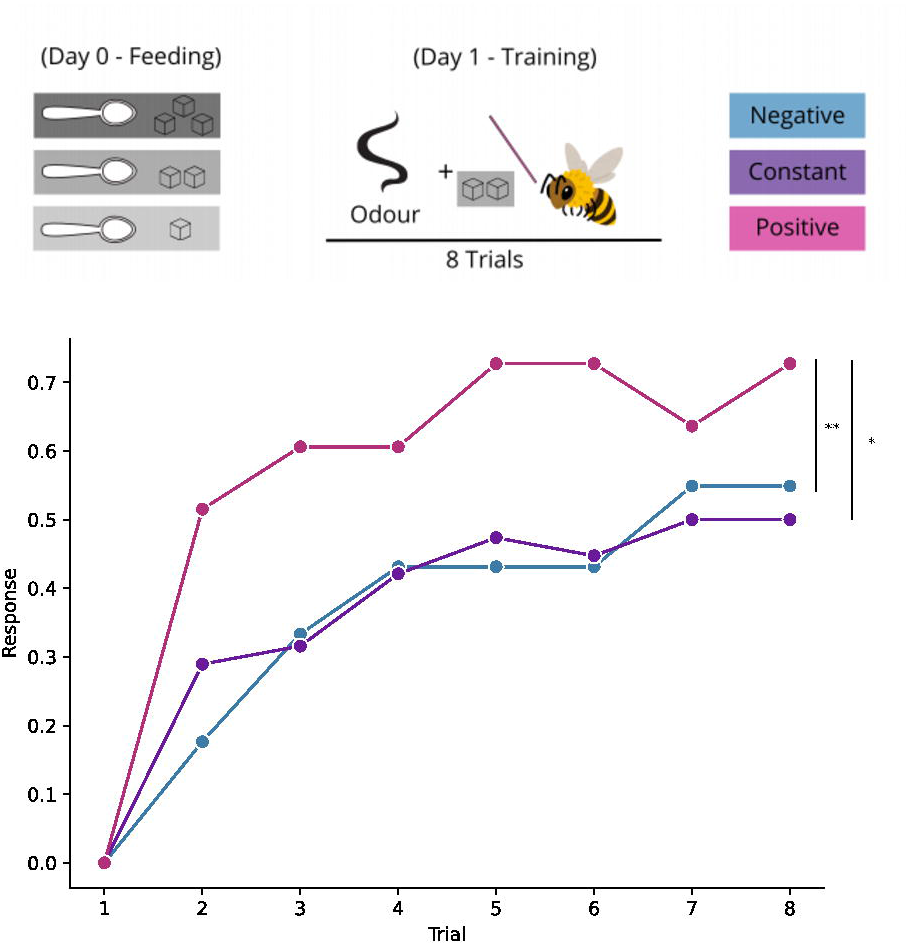
Feeding experience modulates the value of a pre-ingestive reward. (a) Experimental protocol. On day 0 animals were fed with a solution of sucrose 0.1M (light grey), 1M (medium grey) or 1.9M (dark grey) defining three experimental groups: Negative contrast (cyan, n = 51), constant (green, n = 38), and positive contrast (pink, n = 33). On day 1, all animals were stimulated on their antennae with a 1 M sucrose solution, which could be of lower, equal, or higher concentration than the solution received on day 0. (b) line plots showing the proportion of positive responses for each group across the training trials. *p < 0.05, **p < 0.01

**Figure 5.**
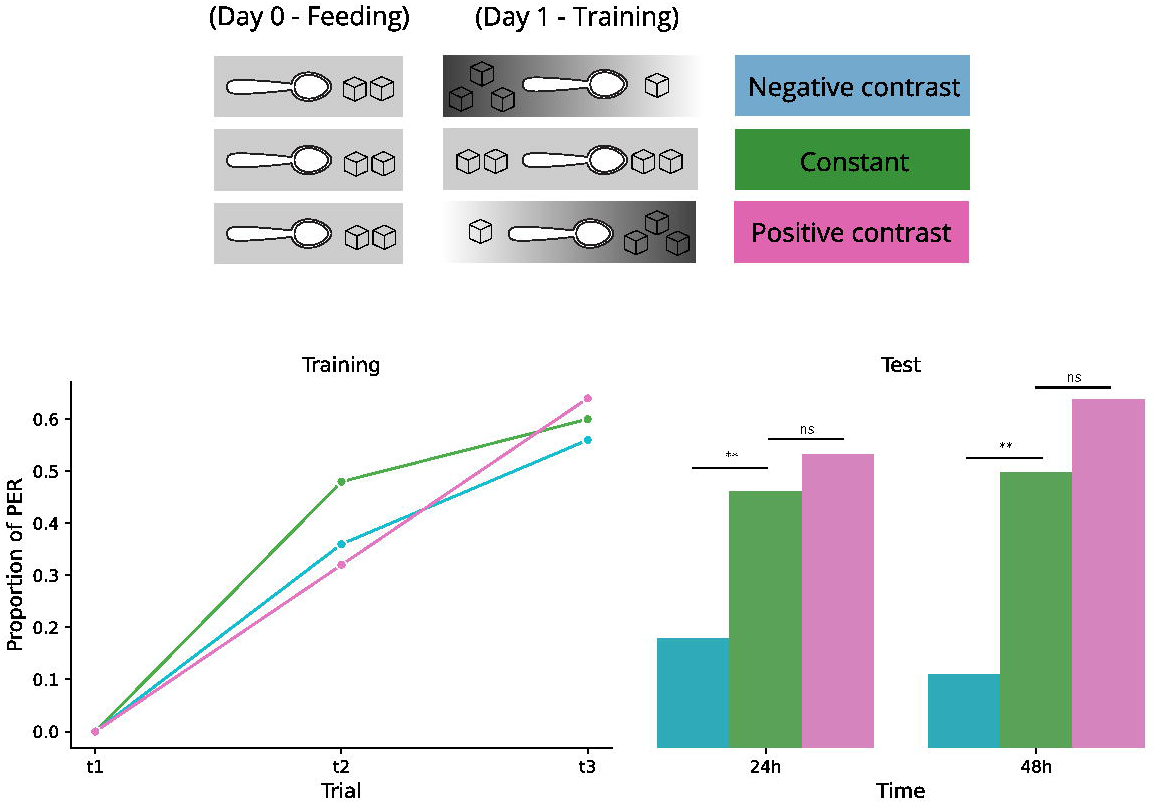
Contrasts between rewards during training modulates memory strength. (a) Experimental protocol. All the animals are fed with a 1M sucrose concentration solution on day 0. On day 1, animals received three trials during training. The reward schedule received during a three trials training defines three experimental groups. Negative contrast (cyan, n = 25), constant (green, n = 25), and positive contrast (pink, n = 25). (b) Left panel: line plots showing the proportion of positive responses for each group across the training trials. Right panel: proportion of positive responses during the tests sessions 24 and 48 hours after training.. ** p < 0.01

**Figure 6.**
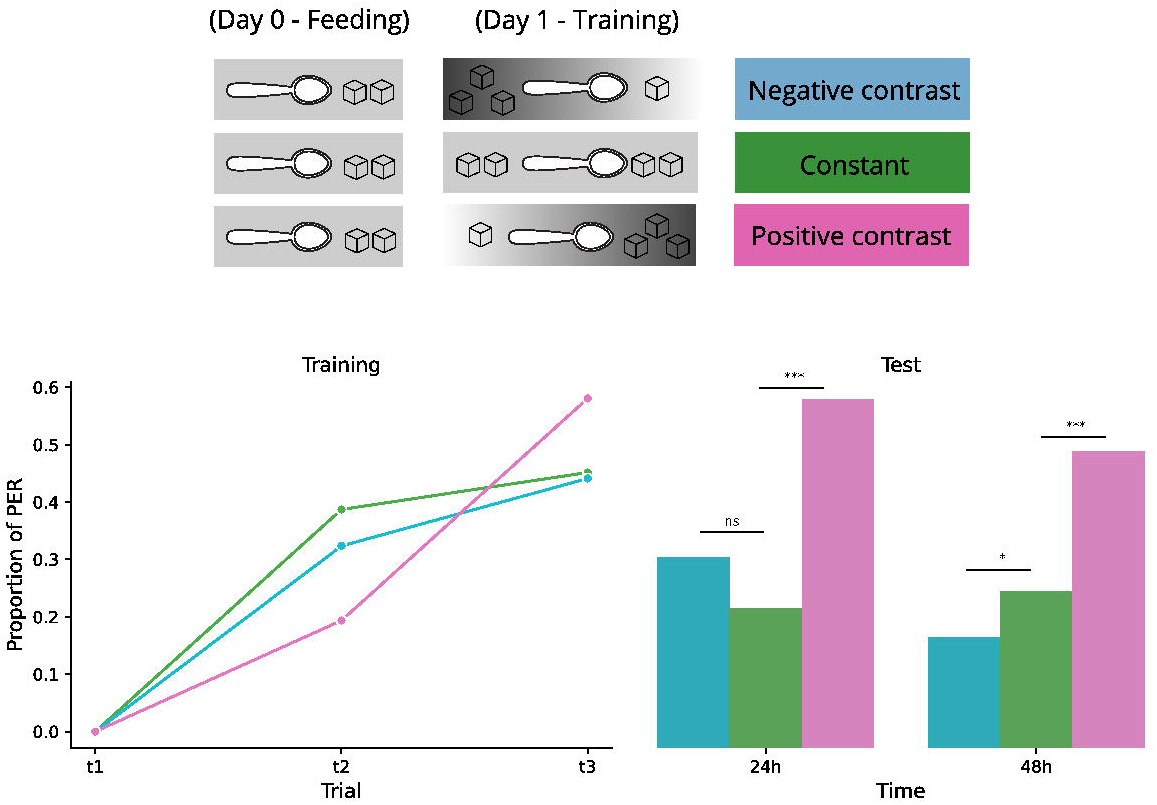
Contrasts in reward between training trials modulate memory strength. (a) Experimental protocol. All the animals were fed with a 0.5M sucrose concentration solution on day 0. The reward schedule received on day 1 defines three experimental groups. Negative contrast (cyan, n = 34), constant (green, n = 31), and positive contrast (pink, n = 31). (b) Left panel: line plots showing the proportion of positive responses for each group across the training trials. Right panel: responses during the tests sessions 24 and 48 hours after training. * p < 0.05, *** p < 0.01

**Figure 7.**
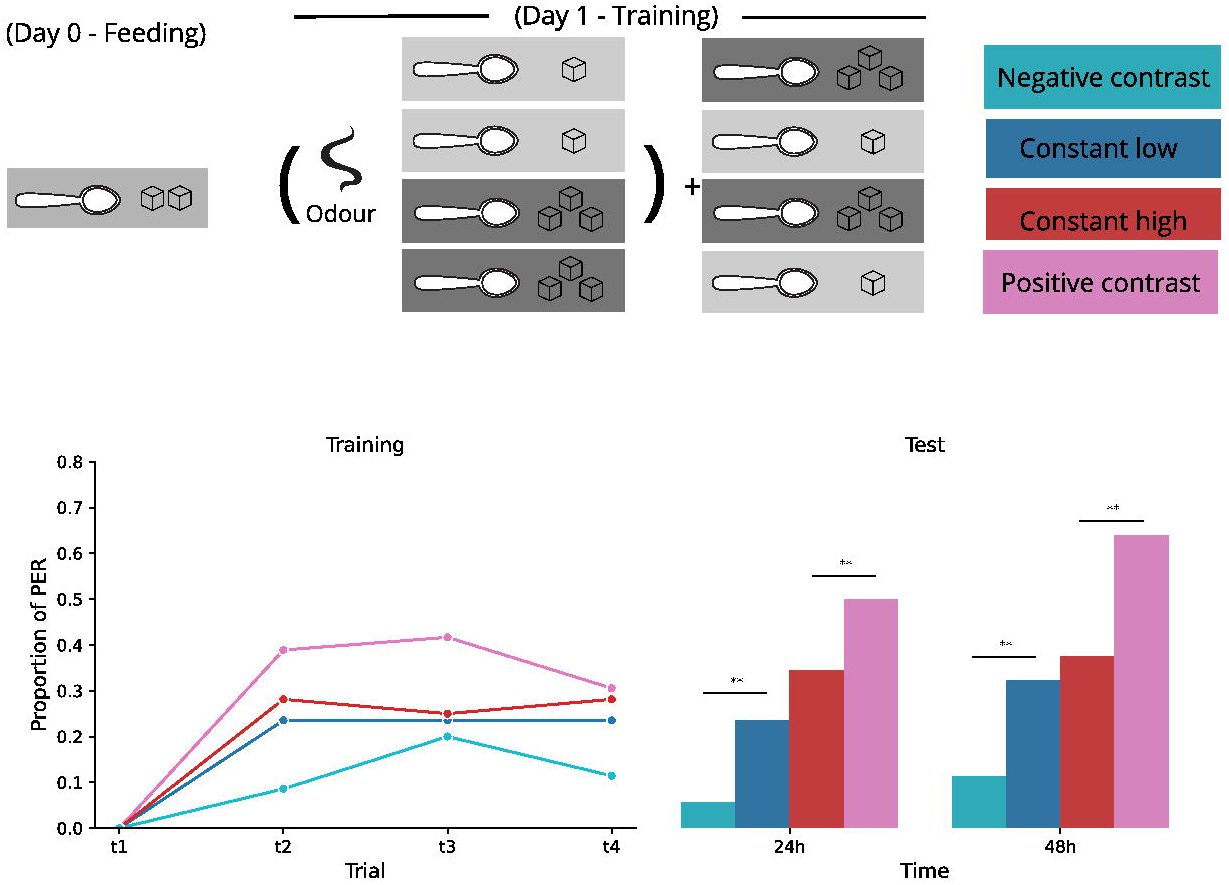
Contrasts between trial and inter-trial rewards modulate memory strength. (a) Experimental protocol. All animals were fed with a 1 M sucrose solution (medium grey) on day 0. On day 1, animals receive a four-trial training with sucrose 0.5M or 1.5M and stimulation between trials with any of the same solutions. The combination of the reward received during training trials and between trials defines four experimental groups. Negative contrast (cyan, n =35), constant low (blue, n =34), constant high (red, n = 32), and positive contrast (pink, n =36). (b) Left panel: line plots showing the proportion of positive responses for each group across the training trials. Right panel: responses during the tests sessions 24 and 48 hours after training. **p < 0.01.

**Figure 8.**
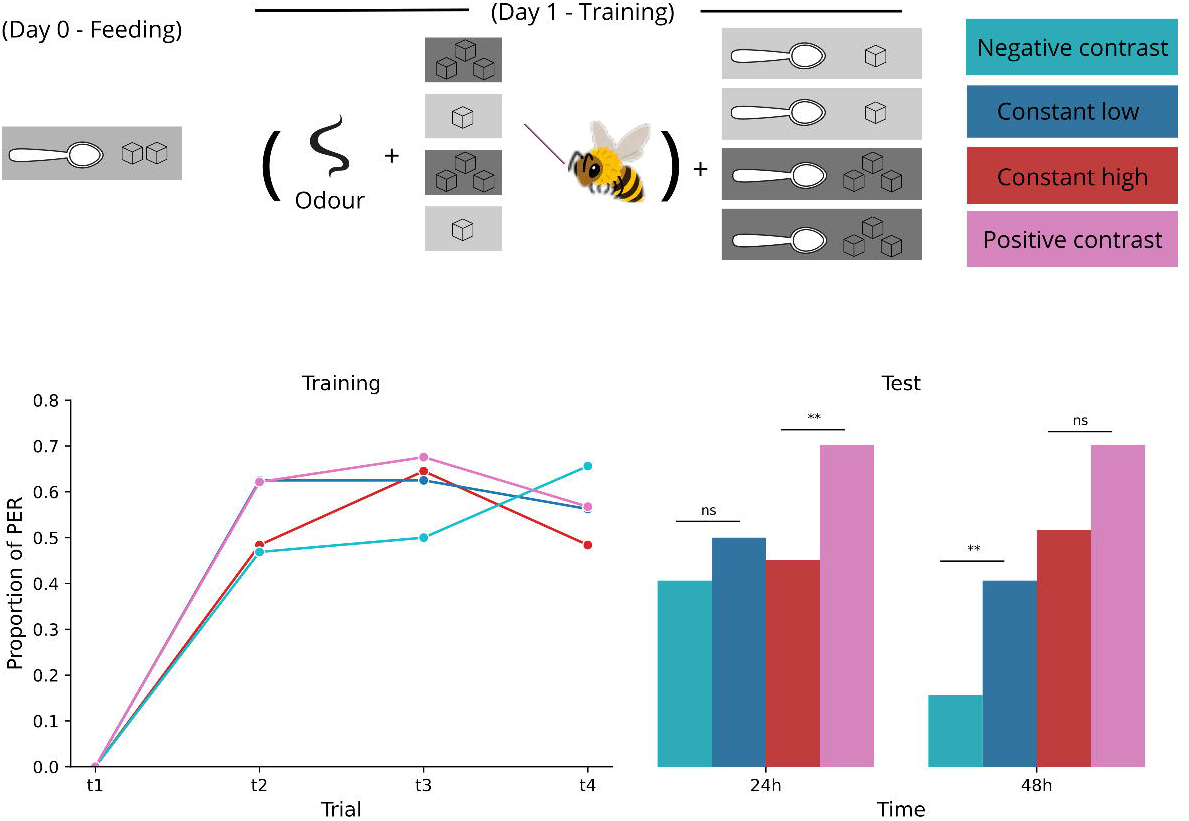
Within-trial reward contrasts modulate memory strength. (a) Experimental protocol. All animals were fed with a 1 M sucrose solution on day 0 (medium grey). On day 1, during each trial, animals were stimulated at their antennae with sucrose 0.5M (light grey) or 1.5M (dark grey) and then fed with a solution of any of those concentrations. The combination of the reward received during training trials and between trials defines four experimental groups. Negative contrast (cyan, n = 33), constant low (blue, n = 32), constant high (red, n =31), and positive contrast (pink, n = 37). (b) Left panel: line plots showing the proportion of positive responses for each group across the training trials. Right panel: responses during the tests sessions 24 and 48 hours after training. *p < 0.05

For analysing the single sensilla recordings, instantaneous spike frequency at 100 ms after the stimulus onset was modeled using Tweedie distribution with a log link function. Experimental group and stimulation solution were included as fixed effects, and individual identity was included as a random intercept to account for repeated measurements. Post hoc pairwise comparisons between groups were performed using estimated marginal means with Tukey correction

Gustatory responsiveness was analysed using a binomial distribution and a logit link function. The response variable was specified as a two-column matrix of successes and failures, corresponding to the number of positive responses and non-responses across six sucrose stimulations. Experimental group was included as a fixed effect, and individual identity was included as a random intercept to account for repeated measurements. The model was fitted using the glmmTMB package in R. Post hoc pairwise comparisons between groups were performed using estimated marginal means with Tukey correction.

## Results

### Feeding experience modulates memory strength 24 hours later

In the standard and widely used honey bee conditioning protocol, animals are captured and harnessed on one day and fed immediately afterward to recover from cold anesthesia and prevent starvation. The conditioning procedure itself begins the following day. This approach is generally preferred because it habituates the bees to restrained conditions and helps control their motivational state during training. Despite the ubiquity of this procedure, the impact of this pre-conditioning feeding regime, and, in particular, the potential contrast between the food provided during this phase and the reward used during training, has not been systematically examined. To address this issue, in the first experiment, we investigated whether food quality on the day preceding a learning session influences memory performance (Fig 1). On day 0, animals were fed either a high (1.5 M) or a low (0.5 M) sucrose concentration solution. Each feeding group was further split into two, resulting in four experimental groups defined by the feeding solution on Day 0 and the reward solution on Day 1. Thus, the groups were “constant high” (fed and trained with 1.5M sucrose solution), “positive contrast” (fed with 0.5M and trained with 1.5M), “constant low” (fed and trained with 0.5M), and “negative contrast” (fed with 1.5M and trained with 0.5M).

We found no significant differences during the acquisition phase between the four groups. By contrast, the relationship between the feeding and reward solutions significantly impacted memory formation. Memory tests performed 24 and 48 hours after training revealed significant differences among groups. The ‘positive contrast’ group exhibited a significantly higher response to the conditioned odour than the ‘constant high’ group (*p* < 0.01), whereas the ‘negative contrast’ group showed a significantly lower response than the ‘constant low’ group (*p* < 0.01). Notably, the differences emerged within the pairs of groups trained with the same reward solution and differing only in the solution ingested on the previous day. We did not find any significant difference between the constant low and constant high groups (p = 0.26). Thus, under these conditions, memory was not modulated by the absolute reward intensity during training but rather by the contrast between the training reward and the solution consumed on the preceding day.

### Sucrose-gustatory responsiveness

Having observed that the quality of food on the day before training modulates memory formation, we asked whether this effect might be due to changes in the perception of reward quality. To address this question, we used a sucrose responsiveness assay that does not involve learning or memory retrieval and therefore provides a direct behavioral proxy for the perceived value of the sucrose solution. Honeybees were collected and fed on day 0 with sucrose solutions of 0.1 M, 1 M, or 1.9 M (Fig 2). These concentrations were chosen to enhance the contrast between treatments. On the following day, we assessed the gustatory responsiveness to a range of sucrose concentrations (R. Scheiner et al., 2001). We observed a significant effect of prior consummatory experience: bees fed a low sucrose concentration on the previous day responded significantly more than those fed medium (p < 0.05) or high sucrose concentrations (p < 0.001), indicating that the quality of the sucrose solution experienced the previous day is sufficient to alter reward evaluation. By contrast, no significant difference was detected between bees fed medium and high sucrose concentrations (p = 0.15). This asymmetry likely reflects the fact that the relative difference between 0.1 M and 1 M sucrose is substantially greater than that between 1 M and 1.9 M.

### Single sensillum recordings

Next, we sought to identify whether the changes in sucrose responsiveness correlate with changes in the sucrose-evoked responses at the level of antennal receptors. We conducted single-sensillum recordings to investigate whether prior experience with sucrose solution modulates the response of gustatory receptor neurons. Using the same three groups of bees described in the previous section, we recorded the responses of chaetic sensilla to different sucrose solutions.

First, we identify the sucrose-sensitive neurons based on their physiological signature. The recorded gustatory receptor neurons (GRNs) exhibited a characteristic phasic response and a concentration-dependent modulation that closely matched the previously described profiles for sucrose receptors in previous works (Haupt 2004).

Interestingly, we found that these sucrose-evoked responses were significantly modulated by the feeding condition on day 0. The instantaneous firing frequency at 100 ms after the stimulus onset revealed distinct sensitivity profiles for the three groups (interaction between stimulus concentration and feeding, p < 0.01). Bees fed a low sucrose concentration (0.1 M) on Day 0 exhibited high sensitivity, reaching a saturation plateau at the lowest stimulation level. In contrast, the medium-concentration group (1 M) showed a reduced initial response that saturated at intermediate stimulation levels. Finally, the group fed the high-concentration (1.9 M) displayed the lowest initial sensitivity, with firing rates that increased progressively across the tested range without reaching a clear plateau.

Collectively, these data suggest a sensory shift in sucrose response profiles, where the dynamic range of the GRNs is calibrated according to the food quality experienced on the previous day. This result indicates that the changes previously observed at the level of behavioural responsiveness and memory formation are accompanied by alterations in the sensitivity of gustatory receptor neurons in the antennae.

### Experience with sucrose modulates pre-ingestive learning

The results from the sucrose responsiveness test and measurements of antennal GRN sensitivity suggest that the modulation of memory induced by the contrast between feeding and reward may, at least in part, arise from changes in sucrose sensitivity. To address this, we used a conditioning protocol in which bees associate an odour with sucrose delivered exclusively to the antennae. This procedure dissociates pre- and post-ingestive components of reward processing, as animals neither ingest the sucrose solution nor contact it with the proboscis. Consequently, no real nutritional feedback is available during training. This type of protocol, in which sucrose is delivered exclusively to the antennae, has been shown to induce short-term, but not long-term memory (Wright et al., 2007). On day 0, we split the animals into three groups, same as in the previous experiment. On day 1, all groups were trained with a 1M sucrose solution as a reward. Our analysis revealed significant differences between groups. We found that the group fed with 0.1M sucrose showed better learning performance than the group fed with 1M (p<0.05) or 1.9M (p<0.01), thus demonstrating that the same concentration of sugar applied to the antennae might represent a stronger or weaker appetitive reward depending on the food consumed 24 hours before.

### Direction of reward change shapes memory formation

In the previous experiments, the differences in learning and memory performance could reflect not only the contrast between feeding and reward conditions, but also to disparities in the total amount of sucrose ingested on Day 0, as bees fed with high- and low-concentration solutions likely differed in their nutritional state. Such differences could themselves influence subsequent learning.To address this limitation and to more closely mimic the temporal dynamics encountered by foraging bees, we designed the following experiment to examine how short-term contrasts in reward quality across successive trials modulate associative memory formation. Importantly, all experimental groups ingested identical total amounts of sucrose on Day 0, as well as during the conditioning protocol on Day 1.

Positive contrast, negative contrast, and no-contrast groups were generated solely by varying the sequence of reward solutions delivered across trials, allowing us to isolate the effect of short-term reward comparisons from metabolic or caloric differences. On Day 0, all bees were fed with a 1 M sucrose solution. On Day 1, they were divided into three groups and subjected to a three-trial conditioning protocol. One group received an ascending reward schedule (0.5–1–1.5 M sucrose), a second group received a constant reward schedule (1 M in all trials), and a third group received a descending reward schedule (1.5–1–0.5 M) (Fig. 5).

At the end of training, all animals had ingested the same total amount of sucrose, ensuring comparable nutritional states both at the beginning and at the end of training.

No significant differences between groups were observed during the training session. We found significant differences between groups during the test sessions 24 and 48 h after training. The ‘constant-’ and the ‘ascending reward’ groups performed better than the ‘descending reward’ group, suggesting that a progressive negative contrast along the conditioning session weakened the formation of appetitive memory. No significant differences were found between the ‘constant reward’ and ‘ascending reward’ groups. We thought that the lack of such an effect possibly reflects a less clear perceptual step between the 1.0 M and 1.5 M sucrose solutions in the last two trials of the ascending protocol.

Therefore, we repeated the same experiment using a geometric progression of reward magnitude. The lowest concentration was 0.17 M sucrose, the intermediate concentration was 0.5 M (threefold higher than the previous one), and the highest concentration was 1.5 M (also threefold higher than the preceding concentration). The constant-reward group received 0.5 M sucrose in all trials (Fig. 6). As in the previous experiment, no significant differences were observed in acquisition performance. During the test session, however, significant differences emerged between the constant-reward and ascending-reward groups, but not between the descending- and constant-reward groups. Despite differences arising from the linear versus geometric progression of reward magnitude, these results indicate that memory formation is sensitive to the direction of fluctuations in reward quality. Importantly, the differences in long-term memory formation observed in this experiment cannot be attributed to modulation of experience strength by nutritional state, but rather to contrasts between expected and experienced reward.

### Unpaired feeding episodes rescale reward value and shape memory

The results obtained so far indicate that the associative strength of a rewarded trial depends on previous experiences with the reward. At least two non-exclusive interpretations arise from these results. First, through odour–reward pairing, honeybees form associations that allow them to predict a specific outcome. Increases or decreases in the quality of this outcome in subsequent trials have opposite effects on memory strength. Second, honeybees may evaluate the quality of a given reward in the context of other available alternatives. In this scenario, an odour that predicts a better option forms a stronger memory than an odour that predicts a poorer one. To distinguish between these possibilities, we conducted the following experiment. On Day 0, bees were collected and fed an intermediate sucrose concentration (1 M). On Day 1, all bees were subjected to a four-trial conditioning session.

Some bees received the conditioned odour paired with a low sucrose concentration (0.5 M) while others received a high sucrose concentration (1.5 M) (Fig. 7). In addition, during the intervals between conditioning trials, all bees received feeding trials without odour stimulation. These feeding episodes had the same volume as the conditioning trials, and the sucrose concentration either matched or differed from that used for conditioning. This design yielded four experimental groups: bees receiving high reward during both conditioning and inter-trial feeding (“constant high”), low reward in both conditions (“constant low”), high reward during conditioning but low reward between trials (“positive contrast”), and low reward during conditioning but high reward between trials (“negative contrast”).

Figure 7 shows performance during training and during the test sessions 24 and 48 h after training. Notably, compared with the previous experiments, both learning rate and memory performance were reduced in this experiment. This was expected, as reward delivery between trials decreases the contingency between the conditioned stimulus and the reward, thereby weakening associative strength. Despite this overall reduction in learning rate, clear differences in memory performance were observed between groups. Consistent with our previous experiments, bees in the “positive contrast” condition exhibited a stronger response to the conditioned odour than bees in the “constant high” condition (p < 0.01), whereas bees in the “negative contrast” condition showed a weaker response than those in the “constant low” condition (p < 0.01). Interestingly, once again, differences in memory were observed between groups that experienced the same sucrose concentration during conditioning trials but differed in the quality of the food provided between trials. Together, these results demonstrate that reward-only episodes delivered between conditioning trials, and not associated with the odor, are sufficient to modulate the perceived value of the reward and subsequent memory formation.

### Contrast between pre- and post-ingestive reward components

In standard PER conditioning protocols, antennal stimulation with sucrose solution is used to elicit the proboscis extension reflex, thereby permitting reward delivery. This sequence parallels the natural foraging process, in which detection of nectar with gustatory receptor neurons in the antenna or tarsa normally precedes proboscis extension and sucrose ingestion. As we observed in previous experiments the antennae are sensitive to different concentrations of sucrose. To investigate whether the memory performance is also modulated by the contrast between the sucrose concentration sensed by the antennae and the sucrose concentration that is ingested, we stimulated the antenna with either a low (0.5M) or high (1.5M) concentration sucrose solution and subsequently rewarded the animals with either a matching or contrasting solution (Fig 8). Animals were divided into four groups based on the sensed and ingested solutions. No significant differences were observed during the training session. Memory test results showed significant differences 24 hs after training between the ‘constant high’ and ‘positive contrast’ groups, but these differences disappear at 48 hs (24hs: p < 0.05, 48hs: p = 0.11). In contrast, between the ‘constant low’ and ‘negative contrast’ groups, no significant differences were found at 24 hs, but at 48 hs significant differences were found between these two groups (24hs: p = 0.43, 48hs: p < 0.05). These results indicate that antennal contact with sucrose alone is sufficient to generate a short-lived expectation, which is hen contrasted with the reward experienced during ingestion.

## Discussion

In this work, we showed that experience modulates how animals respond to subsequent rewards, correlating with changes in the activity of gustatory receptor neurons and exerting a strong influence on learning and memory. We propose that this effect can be understood as the formation of an expectation, where a prior experience modifies the perceived value of a reward. Consequently, the reward is not processed in absolute terms, but rather in relation to whether it is better or worse than expected. Using different experimental protocols, we demonstrate that this effect operates across multiple timescales, ranging from seconds to hours. We do not distinguish here whether this expectation reflects a transient internal state or a stored memory of reward value, as both mechanisms would bias reward evaluation in a similar manner.

Based on our results, we propose that feeding on day 0 generates an internal expectation about reward quality. This experience could trigger the formation of a memory associated with reward evaluation. Supporting this view, a recent study in *Drosophila* showed that preference between different rewards during foraging depends on *rutabaga* expression in the mushroom bodies, consistent with the formation of an expectation-related memory (Martinez-Cordera et al., 2025). In another behavioural task, sucrose presentation produces an internal devaluation when it is encountered in a novel context, suggesting that a single two-minute exposure during learning is sufficient to establish a context-dependent reward expectation (Warnecke et al., 2026). Similarly, in flies, consumption of a non-nutritive sugar induces an internal devaluation of its perceived value. This experience led the authors to propose the formation of a “caloric frustration memory” (CFM), which is accompanied by a reduction in the activity of reward-signalling neurons (Musso et al., 2017).

Our findings show that prior exposure to high sugar concentrations leads to a reduction in gustatory sensitivity. In honeybees, sucrose responsiveness has often been reported to correlate with learning performance in restrained laboratory paradigms (R. Scheiner et al., 2001). However, this relationship does not reflect differences in cognitive ability, but rather variation in reward perception and motivational state, as performance differences disappear when reward magnitude is adjusted to individual responsiveness (Scheiner et al., 2005). Consistent with this view, recent work under free-flying conditions shows that sucrose responsiveness and learning proficiency do not correlate, highlighting the context-dependent nature of this relationship (Kuklovsky et al., 2026).

Although our data demonstrate experience-dependent modulation of gustatory receptor neuron activity, it is expected that reward-related plasticity also occurs at downstream circuit levels. Our findings indicate that changes at the sensory periphery might contribute to bias reward evaluation and, consequently, learning performance, but do not fully account for all experience-dependent effects. In *Drosophila*, exposure to a high-sucrose or high-glucose diet reduces the proboscis extension response (PER) to sugars. Notably, this effect is driven by a decrease in gustatory receptor neuron responsiveness, mediated by alterations in carbohydrate metabolic pathways (May et al., 2019; Sung et al., 2023). Beyond the periphery, both valence-coding neurons—specifically dopaminergic PAM neurons—and mushroom body output neurons (MBONs) also show attenuated responses to sugar following a high-sugar diet, resulting in impairments in associative learning (May et al., 2020; Pardo-Garcia et al., 2023). Together, these findings illustrate how peripheral sensory changes can propagate to central circuits, inducing plasticity that ultimately shapes cognitive performance. In this context, examining how experience with different sugars modulates the activity of VUMmx1, a neuron encoding appetitive reinforcement in honeybees (Hammer, 1993), may provide further insight into the coupling between peripheral and central plasticity.

Our experiments show that positive reward contrasts during training, such as an ascending reward protocol or the availability of food between trials, lead to significantly stronger memory formation. This is consistent with the results reported by Gil and collaborators (Gil et al., 2007), who showed that bees exposed to an artificial feeder delivering increasing reward magnitudes across successive visits exhibit more persistent searching behaviour, whereas a decrease in reward magnitude elicits the opposite response. Dance intensity has likewise been shown to be modulated by the contrast between the reward encountered and prior reward experience (Richter and Waddington, 1993). This pattern has been interpreted as a consequence of reward expectation formation (Gil et al., 2007), an interpretation further supported by similar effects observed in bumblebees and by the reduction in sucrose consumption following a perceived reward downshift (Wiegmann et al., 2003). During foraging, this expectation would allow animals compare the quality of available food across different floral patches, learning to discriminate rewards based on their relative profitability (Hernández et al., 2025; Nityananda and Chittka, 2021). This comparative process operates over short timescales, enabling flexible foraging strategies (Nityananda and Pattrick, 2013). Such flexibility may be particularly advantageous for social insects, as it can promote resource heterogeneity at the colony level. Consistent with this idea, a recent study in bumblebees reported lower flower constancy than previously expected (Yourstone et al., 2023).

In this study, we used pollen foragers in order to minimise behavioural and motivational heterogeneity. Although pollen and nectar foragers are known to differ in their sucrose responsiveness (R Scheiner et al., 2001), these foraging roles are flexible rather than fixed behavioural categories (Arenas et al., 2021). Indeed, individual bees can differ in sucrose responsiveness depending on their immediate foraging context, such as whether they are arriving at or departing from a floral source (Moreno and Arenas, 2024). Our aim was therefore not to characterise traits specific to a particular foraging caste, but to investigate a general cognitive process that is expected to operate across behavioural roles and physiological states.

Our results indicate that identical sensory inputs can be evaluated differently depending on the animal’s recent history, suggesting the involvement of experience-dependent states that shape reward valuation. A conceptually similar mechanism has been described in *Drosophila*, where the selection of oviposition sites has emerged as a tractable model to study decision-making (Yang et al., 2015, 2008)a. These studies showed that females base their decisions not on the absolute properties of individual sites, but on comparisons between available options. More recently, the role of internal expectations in guiding reward-related behaviour has been explicitly demonstrated. Vijayan and collaborators (Vijayan et al., 2022) showed that prior experience establishes an internal expectation that biases egg-laying decisions, such that identical sensory cues are evaluated differently depending on the animal’s history. Building on this work, the same authors proposed a neural model accounting for how such expectation-like signals are implemented (Vijayan et al., 2023). Using an aversive behavioural paradigm, Villar and Pavao et al. showed that the intensity of an electric shock is not evaluated solely in absolute terms, but also relative to the animal’s prior experience (Villar et al., 2022). These studies provide a useful conceptual parallel to our findings. Together, they support the view that experience-dependent internal states rescale the perceived value of rewards before they are incorporated into decision-making or learning processes. Such expectation-like signals may therefore represent a general mechanism through which animals adaptively evaluate rewards or punishment in dynamic environments.

We propose that animals perform reward comparisons across multiple temporal scales. On the one hand, the food received in the laboratory 24 hours before training may generate a long-term expectation, supported by two non-exclusive and potentially causally related mechanisms: a shift in gustatory receptor sensitivity and the formation of a long-term memory associated with reward value. On the other hand, animals also perform short-term comparisons, which may rely on mechanisms similar to those underlying working-memory processes. Together, these mechanisms would allow bees to integrate past and recent experiences to guide efficient foraging decisions in dynamic environments.

## Bibliography

Arenas, A., Lajad, R., Farina, W., 2021. Selective recruitment for pollen and nectar sources in honeybees. J. Exp. Biol. 224, jeb242683. 10.1242/jeb.242683

Bitterman, M.E., Menzel, R., Fietz, A., Schäfer, S., 1983. Classical conditioning of proboscis extension in honeybees (Apis mellifera). J. Comp. Psychol. Wash. DC 1983 97, 107–119.

Couvillon, P.A., Bitterman, M.E., 1984. The overlearning-extinction effect and successive negative contrast in honeybees (Apis mellifera). J. Comp. Psychol. 98, 100–109.

Crespi, L.P., 1942. Quantitative Variation of Incentive and Performance in the White Rat. Am. J. Psychol. 55, 467–517. 10.2307/1417120

De Marco, R.J., Gil, M., Farina, W.M., 2005. Does an increase in reward affect the precision of the encoding of directional information in the honeybee waggle dance? J. Comp. Physiol. A Neuroethol. Sens. Neural. Behav. Physiol. 191, 413–419. 10.1007/s00359-005-0602-3

Gil, M., De Marco, R.J., 2009. Honeybees learn the sign and magnitude of reward variations. J. Exp. Biol. 212, 2830–2834. 10.1242/jeb.032623

Gil, M., De Marco, R.J., Menzel, R., 2007. Learning reward expectations in honeybees. Learn. Mem. Cold Spring Harb. N 14, 491–496. 10.1101/lm.618907

Gil, M., Menzel, R., De Marco, R.J., 2008. Does an insect’s unconditioned response to sucrose reveal expectations of reward? PloS One 3, e2810. 10.1371/journal.pone.0002810

Hammer, M., 1993. An identified neuron mediates the unconditioned stimulus in associative olfactory learning in honeybees. Nature 366, 59–63. 10.1038/366059a0

Haupt, S.S., 2004. Antennal sucrose perception in the honey bee (Apis mellifera L.): behaviour and electrophysiology. J. Comp. Physiol. A Neuroethol. Sens. Neural. Behav. Physiol. 190, 735–745. 10.1007/s00359-004-0532-5

Hemingway, C.T., Muth, F., 2022. Label-based expectations affect incentive contrast effects in bumblebees. Biol. Lett. 18, 20210549. 10.1098/rsbl.2021.0549

Hernández, J.C., García, J.E., Wells, H., Amaya-Márquez, M., 2025. Honey Bee Foraging Decisions Are Shaped by Floral Trait Distinctiveness and Perception of Gains or Losses. Insects 16, 884. 10.3390/insects16090884

Hodgson, E.S., Lettvin, J.Y., Roeder, K.D., 1955. Physiology of a primary chemoreceptor unit. Science 122, 417–418. 10.1126/science.122.3166.417-a

Kuklovsky, V., Avarguès-Weber, A., Giurfa, M., Scheiner, R., 2026. Visual learning performance in free-flying honey bees is independent of sucrose and light responsiveness and depends on training context. Sci. Rep. 16, 1319. 10.1038/s41598-025-34900-9

Marion-Poll, F., 1996. Display and analysis of electrophysiological data under WindowsTM. Entomol. Exp. Appl. 80, 116–119. 10.1111/j.1570-7458.1996.tb00900.x

Martinez-Cordera, M., Sakai, T., Saitoe, M., Ueno, K., 2025. Comparative experience shapes sucrose preference through memory in Drosophila. Mol. Brain 18, 32. 10.1186/s13041-025-01202-0

May, C.E., Rosander, J., Gottfried, J., Dennis, E., Dus, M., 2020. Dietary sugar inhibits satiation by decreasing the central processing of sweet taste. eLife 9, e54530. 10.7554/eLife.54530

May, C.E., Vaziri, A., Lin, Y.Q., Grushko, O., Khabiri, M., Wang, Q.-P., Holme, K.J., Pletcher, S.D., Freddolino, L., Neely, G.G., Dus, M., 2019. High Dietary Sugar Reshapes Sweet Taste to Promote Feeding Behavior in Drosophila melanogaster. Cell Rep. 27, 1675–1685.e7. 10.1016/j.celrep.2019.04.027

Moreno, E., Arenas, A., 2024. Foraging task specialization in honey bees (Apis mellifera): the contribution of floral rewards to the learning performance of pollen and nectar foragers. J. Exp. Biol. 227, jeb246979. 10.1242/jeb.246979

Musso, P.-Y., Lampin-Saint-Amaux, A., Tchenio, P., Preat, T., 2017. Ingestion of artificial sweeteners leads to caloric frustration memory in Drosophila. Nat. Commun. 8, 1803. 10.1038/s41467-017-01989-0

Nityananda, V., Chittka, L., 2021. Different effects of reward value and saliency during bumblebee visual search for multiple rewarding targets. Anim. Cogn. 24, 803–814. 10.1007/s10071-021-01479-3

Nityananda, V., Pattrick, J.G., 2013. Bumblebee visual search for multiple learned target types. J. Exp. Biol. 216, 4154–4160. 10.1242/jeb.085456

Pankiw, T., Waddington, K.D., Page, R.E., 2001. Modulation of sucrose response thresholds in honey bees (Apis mellifera L.): influence of genotype, feeding, and foraging experience. J. Comp. Physiol. [A] 187, 293–301. 10.1007/s003590100201

Pardo-Garcia, T.R., Gu, K., Woerner, R.K.R., Dus, M., 2023. Food memory circuits regulate eating and energy balance. Curr. Biol. CB 33, 215–227.e3. 10.1016/j.cub.2022.11.039

Ramírez, G.P., Martínez, A.S., Fernández, V.M., Corti Bielsa, G., Farina, W.M., 2010. The influence of gustatory and olfactory experiences on responsiveness to reward in the honeybee. PloS One 5, e13498. 10.1371/journal.pone.0013498

Richter, M.R., Waddington, K.D., 1993. Past foraging experience influences honey bee dance behaviour. Anim. Behav. 46, 123–128. 10.1006/anbe.1993.1167

Scheiner, R., Kuritz-Kaiser, A., Menzel, R., Erber, J., 2005. Sensory responsiveness and the effects of equal subjective rewards on tactile learning and memory of honeybees. Learn. Mem. Cold Spring Harb. N 12, 626–635. 10.1101/lm.98105

Scheiner, R., Page, R.E., Erber, J., 2001. Responsiveness to sucrose affects tactile and olfactory learning in preforaging honey bees of two genetic strains. Behav. Brain Res. 120, 67–73.

Scheiner, R, Page, R.E., Jr, Erber, J., 2001. The effects of genotype, foraging role, and sucrose responsiveness on the tactile learning performance of honey bees (Apis mellifera L.). Neurobiol. Learn. Mem. 76, 138–150. 10.1006/nlme.2000.3996

Sung, H., Vaziri, A., Wilinski, D., Woerner, R.K.R., Freddolino, L., Dus, M., 2023. Nutrigenomic regulation of sensory plasticity. eLife 12, e83979. 10.7554/eLife.83979

Tinklepaugh, O.L., 1928. An experimental study of representative factors in monkeys. J. Comp. Psychol. 8, 197–236. 10.1037/h0075798

Vijayan, V., Wang, F., Wang, K., Chakravorty, A., Adachi, A., Akhlaghpour, H., Dickson, B.J., Maimon, G., 2023. A rise-to-threshold process for a relative-value decision. Nature 619, 563–571. 10.1038/s41586-023-06271-6

Vijayan, V., Wang, Z., Chandra, V., Chakravorty, A., Li, R., Sarbanes, S.L., Akhlaghpour, H., Maimon, G., 2022. An internal expectation guides Drosophila egg-laying decisions. Sci. Adv. 8, eabn3852. 10.1126/sciadv.abn3852

Villar, M.E., Pavão-Delgado, M., Amigo, M., Jacob, P.F., Merabet, N., Pinot, A., Perry, S.A., Waddell, S., Perisse, E., 2022. Differential coding of absolute and relative aversive value in the Drosophila brain. Curr. Biol. CB 32, 4576–4592.e5. 10.1016/j.cub.2022.08.058

Warnecke, C., Schweizer, J.A., Zattera, B., Goldschmidt, D., Leptien, K., Felsenberg, J., 2026. Re-exposure to reward re-evaluates related memories. Curr. Biol. CB 36, 565–575.e3. 10.1016/j.cub.2025.11.058

Wendt, S., Strunk, K.S., Heinze, J., Roider, A., Czaczkes, T.J., 2019. Positive and negative incentive contrasts lead to relative value perception in ants. eLife 8, e45450. 10.7554/eLife.45450

Wiegmann, D.D., Wiegmann, D.A., Waldron, F.A., 2003. Effects of a reward downshift on the consummatory behavior and flower choices of bumblebee foragers. Physiol. Behav. 79, 561–566. 10.1016/s0031-9384(03)00122-7

Wright, G.A., Mustard, J.A., Kottcamp, S.M., Smith, B.H., 2007. Olfactory memory formation and the influence of reward pathway during appetitive learning by honey bees. J. Exp. Biol. 210, 4024–4033. 10.1242/jeb.006585

Yang, C.-H., Belawat, P., Hafen, E., Jan, L.Y., Jan, Y.-N., 2008. Drosophila egg-laying site selection as a system to study simple decision-making processes. Science 319, 1679–1683. 10.1126/science.1151842

Yang, C.-H., He, R., Stern, U., 2015. Behavioral and circuit basis of sucrose rejection by Drosophila females in a simple decision-making task. J. Neurosci. Off. J. Soc. Neurosci. 35, 1396–1410. 10.1523/JNEUROSCI.0992-14.2015

Yourstone, J., Varadarajan, V., Olsson, O., 2023. Bumblebee flower constancy and pollen diversity over time. Behav. Ecol. Off. J. Int. Soc. Behav. Ecol. 34, 602–612. 10.1093/beheco/arad028

