## Supplementary material for "Subjective rather than absolute reward value determines long-term memory formation in honey bees": Table 1

| Figure | Phase | Effect |
| --- | --- | --- |
| 1 | TR | Group: χ²=5.62, d.f.=3, P=0.13; Trial: χ²=32.52, d.f.=2, P<0.001; Trial x Group: χ²=5.41, d.f.=6, P=0.49 |
| 1 | TS | Group: χ²=16.82, d.f.=3, P<0.001; Trial: χ²=0.68, d.f.=2, P=0.41; Trial x Group: χ²=2.07, d.f.=3, P=0.56 |
| 2 |  | χ²= 16.70, d.f.=2, P<0.001 |
| 3 |  | Group: χ²= 4.96, d.f.=2, P<0.84; Concentration: χ²=11.69, d.f.=2, P<0.003; Trial x Group: χ²=16.00, d.f.=4, P=0.003 |
| 4 |  | Group: χ²= 9.01, d.f.=2, P<0.05; Trial: χ²=44.61, d.f.=6, P<0.001; Trial x Group: χ²=12.02, d.f.=12, P=0.44 |
| 5 | TR | Group: χ²=0.48, d.f.=2, P=0.79; Trial: χ²=7.68, d.f.=1, P<0.01; Trial x Group: χ²=1.73, d.f.=2, P=0.42 |
| 5 | TS | Group: χ²=8.04, d.f.=2, P=<0.05; Trial: χ²=0.18, d.f.=1, P=0.67; Trial x Group: χ²=1.97, d.f.=2, P=0.37 |
| 6 | TR | Group: χ²=0.01, d.f.=2, P=0.96; Trial: χ²=5.04, d.f.=1, P<0.05; Trial x Group: χ²=5.45, d.f.=2, P=0.07 |
| 6 | TS | Group: χ²=32.52, d.f.=2, P<0.001; Trial: χ²=2.88, d.f.=1, P=0.09; Trial x Group: χ²=12.60, d.f.=2, P<0.01 |
| 7 | TR | Group: χ²= 4.15, d.f.=3, P=0.25; Trial: χ²=1.20, d.f.=2, P=0.55; Trial x Group: χ²=5.30, d.f.=6, P=0.51 |
| 7 | TS | Group: χ²=16.20, d.f.=3, P<0.01; Trial: χ²=3.49, d.f.=1, P=0.06; Trial x Group: χ²=0.81, d.f.=3, P=0.85 |
| 8 | TR | Group: χ²= 0.97, d.f.=3, P=0.81; Trial: χ²=1.87, d.f.=2, P=0.39; Trial x Group: χ²=9.49, d.f.=6, P=0.15 |
| 8 | TS | Group: χ²= 10.76, d.f.=3, P<0.05; Trial: χ²=1.47 d.f.=1, P=0.23; Trial x Group: χ²=5.85, d.f.=3, P=0.12 |
